# Rapid *in vivo* development of resistance to daptomycin in vancomycin-resistant *Enterococcus faecium* due to genomic rearrangements

**DOI:** 10.1101/2021.09.24.461763

**Authors:** Sarah Mollerup, Christine Elmeskov, Heidi Gumpert, Mette Pinholt, Tobias Steen Sejersen, Martin Schou Pedersen, Peder Worning, Dorte Frees, Henrik Westh

**Author notes:** Corresponding author: Sarah Mollerup, PhD, Department of Clinical Microbiology, Hvidovre Hospital, Kettegård Alle 30, DK2650 Hvidovre, Denmark.

## Abstract

**Background:** Daptomycin is a cyclic lipopeptide used in the treatment of vancomycin-resistant *Enterococcus faecium* (VREfm). However, the development of daptomycin-resistant VREfm challenges the treatment of nosocomial VREfm infections. Resistance mechanisms of daptomycin are not fully understood. Here we analysed the genomic changes leading to a daptomycin-susceptible VREfm isolate becoming resistant after 40 days of daptomycin and linezolid combination therapy.

**Methods:** The two isogenic VREfm isolates (daptomycin-susceptible and daptomycin-resistant) were analysed using whole genome sequencing with Illumina and Nanopore.

**Results:** Whole genome comparative analysis identified the loss of a 46.5 kb fragment and duplication of a 29.7 kb fragment in the daptomycin-resistant isolate, with many implicated genes involved in cell wall synthesis. Two plasmids of the daptomycin-susceptible isolate were also found integrated in the chromosome of the resistant isolate. One nonsynonymous SNP in the *rpoC* gene was identified in the daptomycin-resistant isolate.

**Conclusions:** Daptomycin resistance developed through chromosomal rearrangements leading to altered cell wall structure. Such novel types of resistance mechanisms can only be identified by comparing closed genomes of isogenic isolates.

## Introduction

Enterococci are commonly found in the environment, in the human and animal gut as commensal bacteria and as hospital-adapted pathogens(1). *Enterococcus faecium* is an important pathogen in nosocomial infections, where it causes a variety of infections such as urinary tract infections, intraabdominal infections, catheter related infections and bacteremia(1). Vancomycin is first-line treatment of *E. faecium* infections, however the emergence of vancomycin-resistant *E. faecium* (VREfm) has limited the treatment options(2, 3). VRE treatment is often linezolid or daptomycin sometimes given in combination(4). Daptomycin is a bactericidal cyclic lipopeptide that disrupts multiple bacterial cell membrane functions(5).

Whole genome sequencing (WGS) of *E. faecium* isolates has led to the identification of several chromosomal loci associated with decreased daptomycin susceptibility (6). Frequently, the identified genetic changes map into genes that can be divided into two functional categories: (i) genes encoding regulatory systems responding to cell envelope stress such as the *liaFSR*, and the *yycFG/yycHIJ* genes encoding homologues of two- or three-component systems that in other bacteria respond to antibiotics targeting the cell wall, and, (ii) genes encoding enzymes involved in the metabolism of cell membrane phospholipids such as *cls*, a cardiolipin synthase, *GdpD*, a glycerolphosphoryl diester phosphdietersterase, and *mprF*, a multipeptide resistance factor. Other studies have failed to identify mutations in the abovementioned genes in daptomycin-resistant *E. faecium* strains(7), have identified other mutations(8), or have also detected presumed resistance mutations prior to daptomycin treatment (9). The mechanisms of daptomycin resistance thus seems to be diverse.

We applied WGS to VREfm isolates from a patient obtained before and after development of daptomycin resistance. The aim of the study was to investigate the genetic background leading to the development of daptomycin resistance in VREfm following prolonged daptomycin and linezolid combination therapy.

## Methods

### Patient history

The patient was admitted to the intensive care unit of Copenhagen University Hospital Hvidovre (Copenhagen, Denmark) with gallstone induced pancreatitis, which developed into necrotising pancreatitis with intraabdominal abscesses. The patient was treated with vancomycin and broad-spectrum antibiotics, and the abscesses were flushed with vancomycin. After thirty days, a VREfm isolate was identified (minimum inhibitory concentration (MIC) of daptomycin 2 mg/l), and combination therapy with linezolid and daptomycin was started. Forty days later, the patient had developed daptomycin-resistant VREfm (MIC 8 mg/l), and treatment was changed to linezolid (MIC 0.5 mg/l) and tigecycline (MIC 0.016 mg/l).

### Isolates

The daptomycin susceptible VREfm isolate, V1164, was identified in drain fluid from an intraabdominal abscess, and the first daptomycin-resistant VREfm isolate, V1225, was identified from the patient’s bloodstream. Vancomycin, linezolid, tigecycline, and rifampicin MICs were established using E-test and applying the EUCAST breakpoints, while an ECOFF value for *E. faecium* of ≤4 μg/ml was applied for daptomycin(10).

### Whole genome sequencing

WGS of the two isolates was performed using an Illumina MiSeq using Nextera XT library preparation kit running 2×150 bp paired-end reads as previously described(11). High molecular weight DNA was obtained with the GenFind V2 kit (Beckman Coulter, Brea, USA) from 3 ml SB overnight cultures and Oxford Nanopore sequencing was performed with a SQK-LSK108 1D ligation kit and native barcoding indices on a R9.4.1 flow cell

### Bioinformatic analysis

Nanopore reads were basecalled using Albacore software and barcodes were trimmed using porechop with an additional demulitplexing check compared to the Albacore barcode splitting. A hybrid assembly using Nanopore and Illumina reads was created using Unicycler v. 4.0.7 with default settings(12). Nanofilt(13) was used to filter nanopore reads. Application of different cut-offs for quality filtering scores prior to hybrid assembly were tested. For V1225, the final closed assembly was obtained removing reads with quality score <10. Short, linear contigs were discarded from the final assemblies.

Core-genome and multi-locus sequencing types (cgMLST and MLST, respectively) were assigned using Ridom SeqSphere+ (Ridom GmbH). Vancomycin-resistance genes were detected using an in-house script. The genomes were annotated using Prokka(14) v. 1.14.5. The NASP single nucleotide polymorphism (SNP) pipeline was used to identify SNPs in V1225 using the closed V1164 chromosome as reference. This included masking of duplicate regions using NUCmer(15), mapping of reads using BWA(16), and SNP calling and identification using GATK(17) with default thresholds. The effect of identified SNPs were evaluated using web-based BLASTx of of the gene sequences containing identified mutations.

Genomic rearrangements were identified by aligning the two closed genomes using MUMmer(18) v. 4.0.0beta2 setting the minimum length of a match (-l) to 3000, computing forward and reverse complement matches (-b), and including non-unique matches in the reference sequence (-maxmatch). The rearrangements were visualized using Ribbon(19).

To confirm loss or acquisition of genetic information, Illumina reads were aligned against the closed genomes using Bowtie 2(20) v. 2.3.4.1 (adding options --no-mixed -X 2000). Duplicate reads were removed using Picard toolkit v. 2.20.2 MarkDuplicates function(21), depth and breadth of coverage was assessed using BEDtools v. 2.28.0(22), and genome coverage was plotted in R v. 4.0.0(23). Command-line analysis jobs were executed in parallel using GNU parallel v. 20181222(24).

The raw reads are available from the European Nucleotide Archive (V1164 Sample accession SAMEA5367917, V1225 Sample accession SAMEA5367970) and the two closed genomes are deposited in GenBank under accession numbers CP083912-CP083929.

### Transmission electron microscopy

The two isolates V1164 and V1225 were grown overnight at 37°C on brain heart infusion agar plates. The next day the cultures were diluted 1:10 in 10 ml BHI medium grown at 37°C with shaking (200 rpm) until OD_600_ reached approximately 1.0. When OD_600_ was reached, the cultures were put on ice until imaging. A negative stain procedure single-droplet method was used before imaging; formvar-carbon coated grids (Pure C, 200 mesh Cu) were glow discharged (30 sec., 5 mA) before use to increase their hydrophilicity. Then 10 μl of the sample was placed on the grid. After 60 seconds the excess sample was slowly removed from the opposite side using a wedge of filter paper and10 μl of staining (2% phosphotungstic acid, pH = 7) was applied. After additionally 60 seconds excess staining from the opposite side was removed using a wedge of filter paper and a rinse with distilled water was performed. Subsequently samples were examined with a Philips CM 100 Transmission EM™ (Philips, Eindhoven, The Netherlands), operated at an accelerating voltage of 80 kV. Images were recorded with an OSIS Veleta™ digital slow scan 2,000 × 2,000 CCD camera and the iTEM™ (Philips) software package.

## Results

The daptomycin-susceptible isolate V1164 exhibited a daptomycin MIC of 2 mg/l and a rifampicin MIC of 8 mg/l while the daptomycin-resistant isolate V1225 exhibited a daptomycin MIC of 8 mg/l and a rifampicin MIC of ≥32 mg/l.

### Genetic events leading to daptomycin resistance in VREfm V1225

WGS of the isolates was performed to investigate their genetic background and to uncover potential mutations causing daptomycin resistance. Hybrid assembly of Illumina and Nanopore reads resulted in closed genomes of both the daptomycin-susceptible V1164 and the daptomycin-resistant V1225 (Supplementary table 1). The two VREfm isolates were assigned to cgMLST 864, MLST 18, CC17. The *vanA* gene was present on a circular plasmid of 52 kb in V1664 and of 48 kb in V1225. One SNP had developed in the daptomycin-resistant VREfm isolate V1225 compared to the susceptible V1164. The identified A->G substitution in position 3485 was a missense mutation in *rpoC* encoding the DNA-directed RNA polymerase subunit beta’, resulting in the amino acid change K1163E (lys->glu).

Whole genome comparative analysis (Figure 1) showed that a 46.5 kb region in the closed chromosome of V1164 (Supplementary file 1) was absent in the V1225 genome. The deleted region contained 45 genes (Supplementary table 2). Some of the absent genes were transposases, genes encoding capsular biosynthesis proteins, genes associated with mannose metabolic pathways and gene encoding a chloride channel protein, but none have previously been linked to daptomycin resistance. Immediately downstream of the deletion, an insertion of a duplicated 29.7 kb region (Supplementary file 2) containing 32 genes was identified (Supplementary table 3). These included genes involved in iron homeostasis. Two plasmids of 21.1 kb and 3.9 kb present in V1164 were found to be integrated in the chromosome of V1225, these comprised 15 and 5 genes, respectively (Supplementary table 4 and 5) and included genes involved in translocation of substrates across membranes. ISL3 family transposase genes bordered all the affected genomic regions, suggesting that these genes may have been involved in the recombination events.

**Figure 1.**
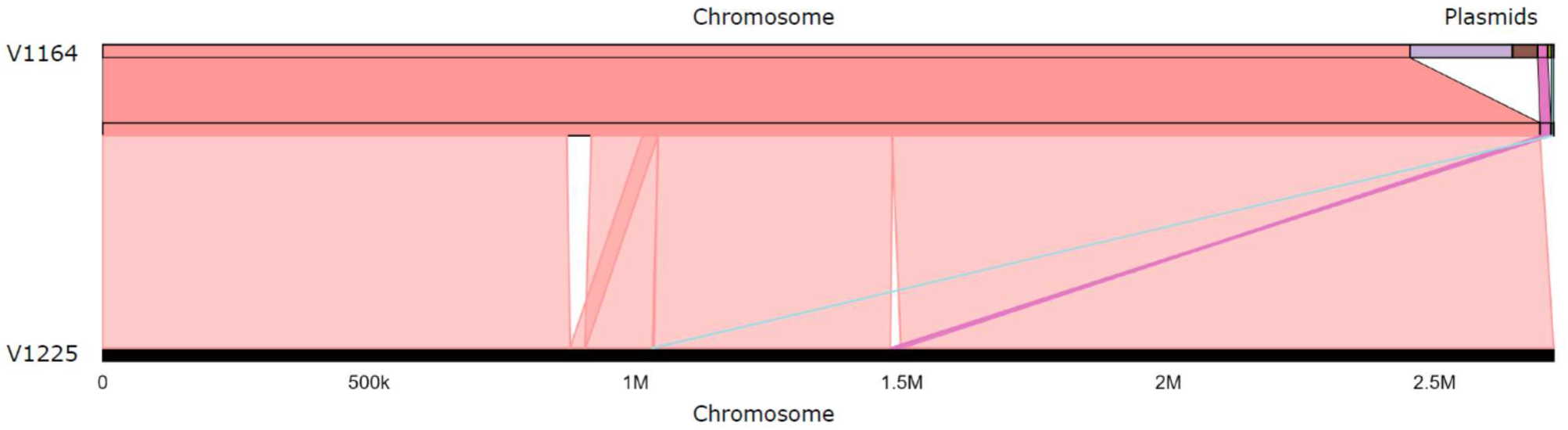
Genome rearrangements in the daptomycin-resistant VRE isolate. Visualization of alignment of the daptomycin-susceptible (V1164) and the daptomycin-resistant (V1225) VRE isolates. The top part represents the V1164 genome (chromosome and plasmids), and the bottom part represents the V1225 chromosome. The cyan and magenta lines indicate insertion of two V1164 plasmids in the V12225 chromosome.

To confirm the deletion and duplication events, reads were mapped to the closed genomes. Mapping of V1225 reads to the V1164 chromosome showed missing coverage at the chromosomal deletion and a depth of coverage of twice the average depth at the chromosomal region being duplicated in V1225 (Supplementary Figure 1). Mapping of V1664 reads to the V1225 chromosome showed a depth of coverage of half the average depth at the duplicated region (and the original location), confirming these genomic rearrangements. Integration of the two plasmids in V1225 could not be confirmed by mapping, due to ISL3 family transposases being present both in the plasmids and at the integration sites, hence the sequence representing the transition from plasmid to chromosome was already present and no gap in coverage could be observed. V1164 was Illumina-sequenced twice, and hybrid assembly using the second set of read pairs also assembled the 21.1 kb plasmid, in this case with higher depth of coverage than obtained for the first hybrid assembly, indicating that the sequence does represent a plasmid (Supplementary table 6). Application of different levels of quality filtering of the nanopore reads also resulted in “integration” of this plasmid into the chromosome, indicating that a mixed population could exist for V1164 with some bacterial cells having the plasmid integrated and some not.

### Electron microscopy of the daptomycin-susceptible and daptomycin-resistant isolate

Previous studies demonstrated that the development of daptomycin resistance in *E. faecalis* is associated with profound ultrastructural changes in the cell envelope(25). This prompted us to perform electron microscopy of both the susceptible and resistant isolates to investigate any morphological changes between the two strains. Transmission electron microscopy (TEM) revealed notable differences in the cell morphology of the two isolates (Figure 2). The daptomycin-resistant VREfm V1225 cells appeared larger than the susceptible V1164 strain, and while the V1164 strain grew in chains, V1225 cells tended to clump and form aggregates. Moreover, characteristic abrasions were visible in the cell envelope of V1225 cells possibly leading to budding (see arrows in Figure 2).

**Figure 2.**
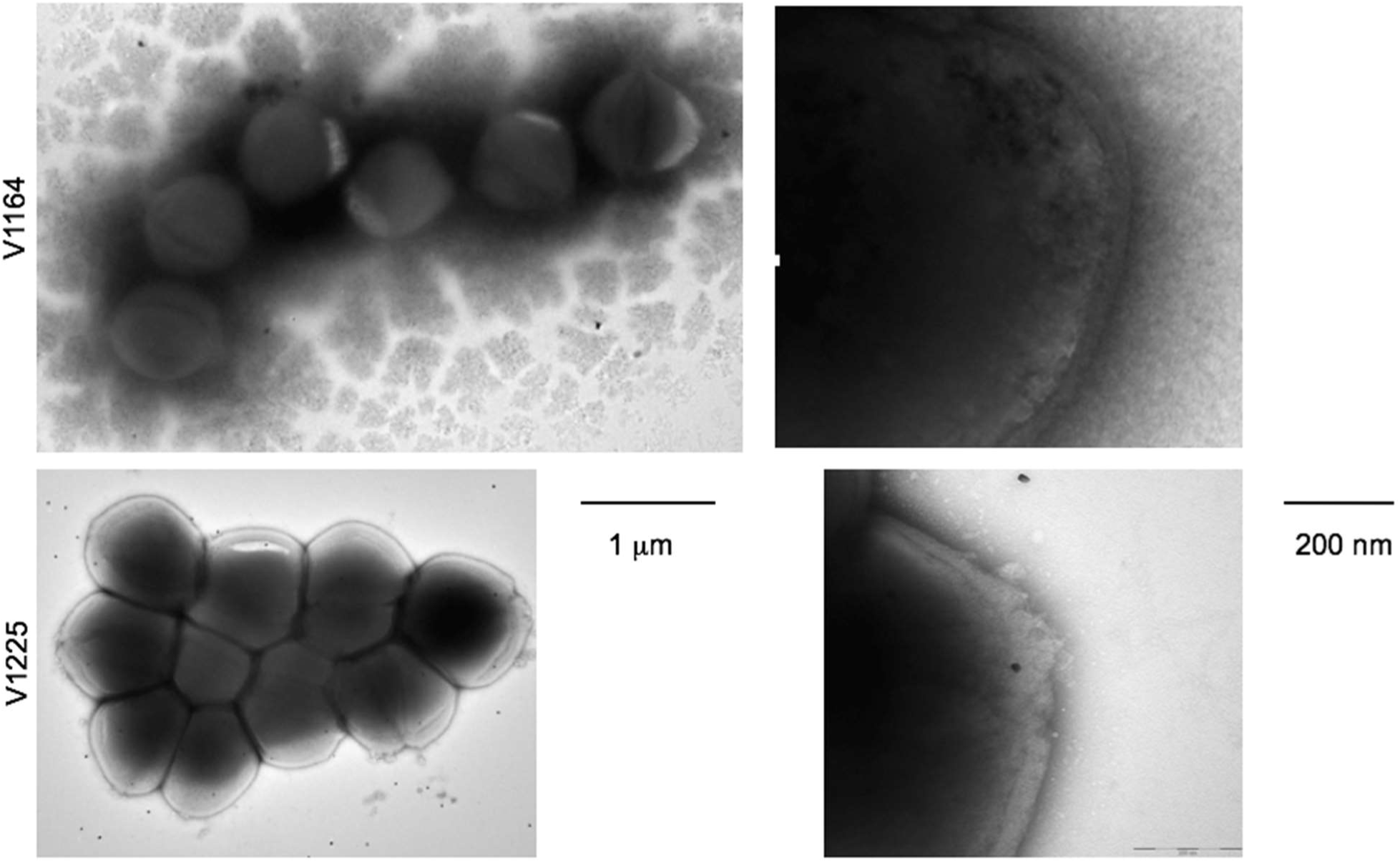
Electron microscopy of the two VRE isolates. Negative stain electron microscopy of daptomycin-susceptible V1164 (top) and daptomycin-resistant VRE V1225 (bottom). Arrows point to abrasions in the cell wall with white vacuoles released from the abrasion.

## Discussion

Daptomycin resistance in enterococci is still a rare phenomenon, but it is of great concern for the individual patient(6). Daptomycin is mainly excreted by the kidney, but a low amount is also excreted in the faeces. As VREfm colonization is localized to the gut, the low gut concentration of daptomycin might contribute to the development of resistance in VREfm(26).

The 46 kb loss of chromosomal genes could be the cause of daptomycin resistance, possibly in combination with the 29 kb duplication. However, the duplication and insertions could also be part of compensatory genomic mechanisms to reduce the impact of the deleted genes. Interestingly the loss involved genes annotated as involved in the mannose metabolic pathway presumably decreasing glycolisation of cell wall structural proteins, and genes of the capsular polysaccharide biosynthesis pathway(27). The loss of these genes may be the cause of the altered cell and cell wall structure seen by TEM. The large deletion found is reminiscent of a finding where a *Pseudomonas aeruginosa* isolate lost 8% of its genome following repeated antimicrobial courses over years of treatment in a patient with cystic fibrosis(28). The duplication of genes involved in iron homeostasis could be a compensatory mechanism in response to the host iron withholding defense system limiting the availability of free iron(29).

Several genetic mutations, predominantly SNPs and often in combination, have been associated with daptomycin resistance in VREfm(6), but no specific mutation has been pinpointed as the leading cause of resistance. We identified one SNP in the *rpoC* gene of the daptomycin-resistant VREfm isolate V1225. Mutations in *rpoB* and *rpoC* have previously been identified in *Staphylococcus aureus* strains exhibiting increased MICs for daptomycin(30, 31) and rifampicin(31), and the here identified mutation could therefore contribute to the decreased susceptibility to both daptomycin and rifampicin observed.

The daptomycin-resistant VREfm V1225 isolate exhibited a number of morphological changes such as increased cell size, cell clumping, and characteristic distortions in the cell envelope. Interestingly, similar morphological changes were previously observed in daptomycin-resistant enterococci that have become resistant through mutations in other genes (*gdpP, liaF*, and *cls*)(25). Release of membrane phospholipids that inactivate daptomycin has been reported for Staphylococci, Streptococci, and *E. faecalis*, thus representing another mechanism leading to daptomycin treatment failure. Our isolates ability to rapidly mutate over weeks reflects the adaptability of the enterococcal genome when challenged by continuous antimicrobial treatment. Interestingly, concomitant treatment with linezolid did not prevent development of daptomycin resistance. In conclusion, our daptomycin-resistant VREfm had a large chromosomal deletion, a duplication and two plasmid insertions leading to an altered cell structure. These genetic events would have been unrecognized by antimicrobial resistance detection software and were only identified because closed genomes of the daptomycin-resistant isolate and its isogenic ancestor were compared.

## Acknowledgements

We would like to express our gratitude to The Core Facility for Integrated Microscopy (http://cfim.ku.dk) for support with electron microscopy. Computerome 2.0 was used for computational analysis.

## Transparency declarations

None to declare

## Supplementary figures

**Supplementary figure 1.**
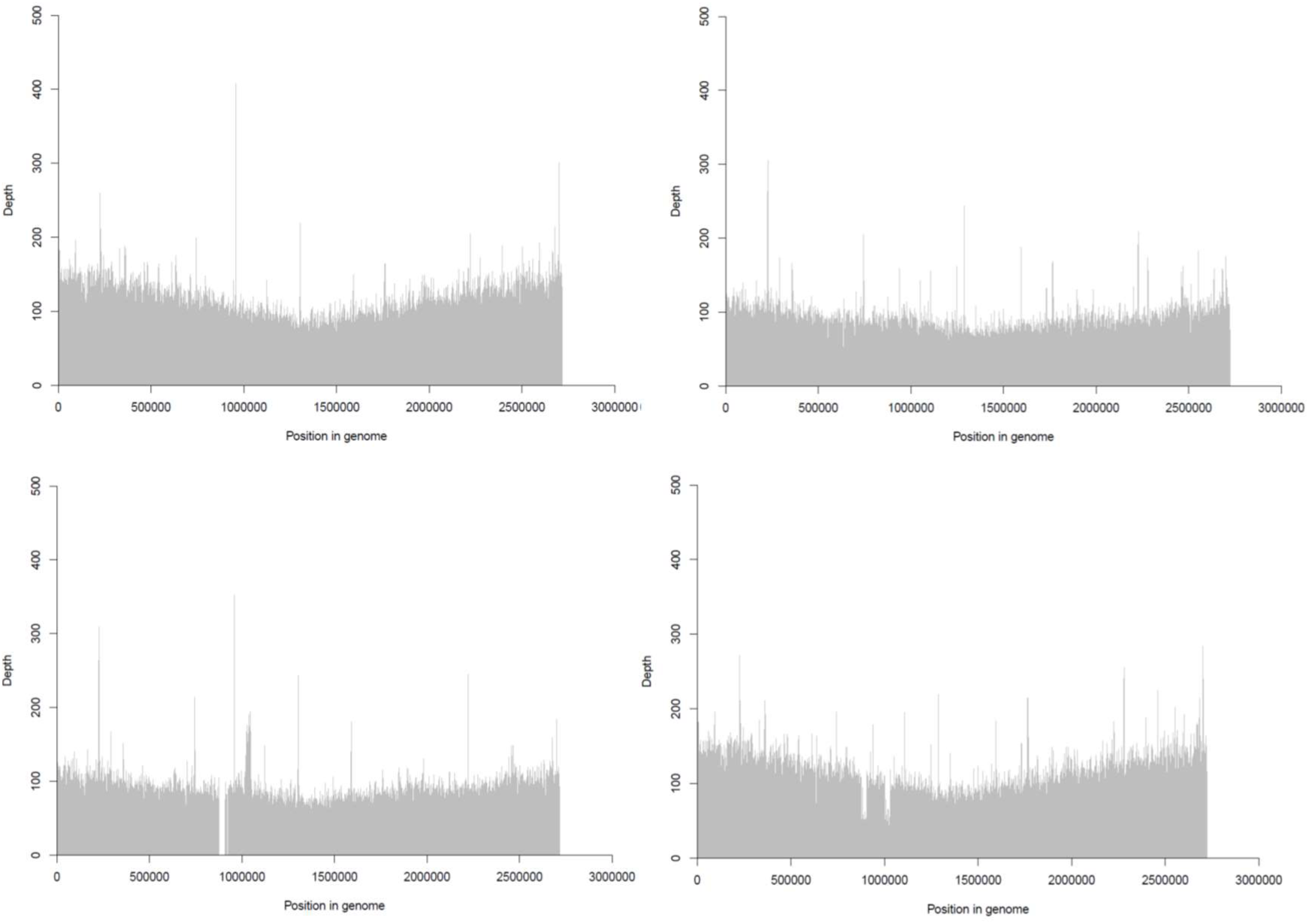
Genome coverage of the two VRE isolates. Top left: V1164 reads mapped to V1164 chromosome; top right: V1225 reads mapped to V1225 chromosome; bottom left: V1225 reads mapped to V1164 chromosome; bottom right: V1164 reads mapped to V1225 chromosome.

